# Brain dopamine imbalance causes follicle death and underlies negative effect of high sugar diet during *Drosophila* oogenesis

**DOI:** 10.1101/2025.04.25.650701

**Authors:** Rodrigo Dutra Nunes, Daniela Drummond-Barbosa

## Abstract

Unhealthy diets, obesity, and low fertility are associated in *Drosophila* and humans. We previously showed that a high sugar diet, but not obesity, reduces *Drosophila* female fertility owing to increased death of newly formed germline cysts and vitellogenic follicles. *Drosophila* strains carrying mutations in the *yellow* (*y*) and *white* (*w*) pigmentation genes are routinely used for investigating the effects of high sugar diets, but it has remained unclear how this genetic background interacts with high sugar. Here, we show that wildtype females retain normal fertility on high sugar compared to control diets, and that mutation of *y* is responsible for the previously observed vitellogenic follicle death on high sugar. The known requirement of *y* for melanin biosynthesis from dopamine, as well as the association between high sugar consumption and reduced dopamine in mammals and decreased dopamine responses in male *Drosophila*, prompted us to investigate potential connections between *y*, high sugar, dopamine and oogenesis. We found that global impairment of dopamine metabolism leads to vitellogenic follicle degeneration while alleviating dopamine imbalance in *y* mutant females prevents follicle death on a high sugar diet. Finally, lack of dopamine production in the central nervous system is sufficient for vitellogenic follicle death on a high sugar diet, and severe dopamine imbalance causes follicle death regardless of diet or genetic background. Our findings are broadly relevant to our understanding of how the effects of unhealthy diets might differ depending on genetic factors and highlight a key connection between brain dopamine metabolism and ovarian follicle survival.

**ARTICLE SUMMARY:** Unhealthy diets, obesity, and reduced fertility are associated in *Drosophila* and humans. Recently, we showed that a high sugar diet induces ovarian follicle death and reduces *Drosophila* fertility independently of obesity. Here, we report that follicle degeneration induced by a high sugar diet depends on genetic background in connection with brain dopamine imbalance. We also show that severe dopamine imbalance increases follicle death regardless of diet or genetic background. These findings are broadly relevant to our understanding of how the effects of unhealthy diets might differ depending on genetic factors and highlight an important connection between dopamine metabolism and oogenesis.

## INTRODUCTION

High sugar consumption and obesity are associated with negative health outcomes, including infertility (Stanhope 2016; Faruque et al. 2019; Malik and Hu 2022; Huang et al. 2023). In *Drosophila*, other groups had shown that a high sugar diet leads to obesity and reduced female fertility (Morris et al. 2012; Brookheart et al. 2017), but it had remained unclear if obesity itself had direct negative effects on oogenesis. More recently, we showed that a high sugar diet, but not obesity, causes decreased fertility in *Drosophila* (Nunes and Drummond-Barbosa 2023). However, the underlying mechanisms remained largely unknown.

*Drosophila* is a well-established genetic model for the study of oogenesis processes (Doherty et al. 2022; Spradling et al. 2022; Giedt and Tootle 2023) and has evolutionarily conserved metabolic responses to high sugar diets (Na et al. 2013; Brookheart et al. 2017; May et al. 2020; Sanz et al. 2022; Nunes and Drummond-Barbosa 2023; Yang et al. 2023). Each female has a pair of ovaries subdivided into ∼15 ovarioles with an anterior germarium followed by developing follicles (also known as egg chambers) (Fig. 1A). Follicles, composed of a 16-cell germline cyst (i.e., one oocyte and 15 nurse cells) surrounded by a monolayer of follicle cells, are produced within the germarium. Newly formed follicles continue to develop through 14 discernable stages to form mature oocytes. Once the follicle reaches stage 8, the oocyte begins taking up yolk, marking the onset of vitellogenesis, a resource- and energy-intensive process that is highly regulated in response to physiological changes (Drummond-Barbosa 2019). In particular, we showed that vitellogenic follicles (as well as early germline cysts within the germarium) die at higher rates in females on a high sugar diet compared to those on a control diet (Nunes and Drummond-Barbosa 2023). This increase in follicle death on a high sugar diet is reminiscent of the increased follicle atresia observed in the ovaries of rats maintained on high fat-high sugar diets (de Melo et al. 2021; Rejani et al. 2022; Demirel et al. 2024), suggesting that conserved mechanisms could be at play.

**Fig. 1.**
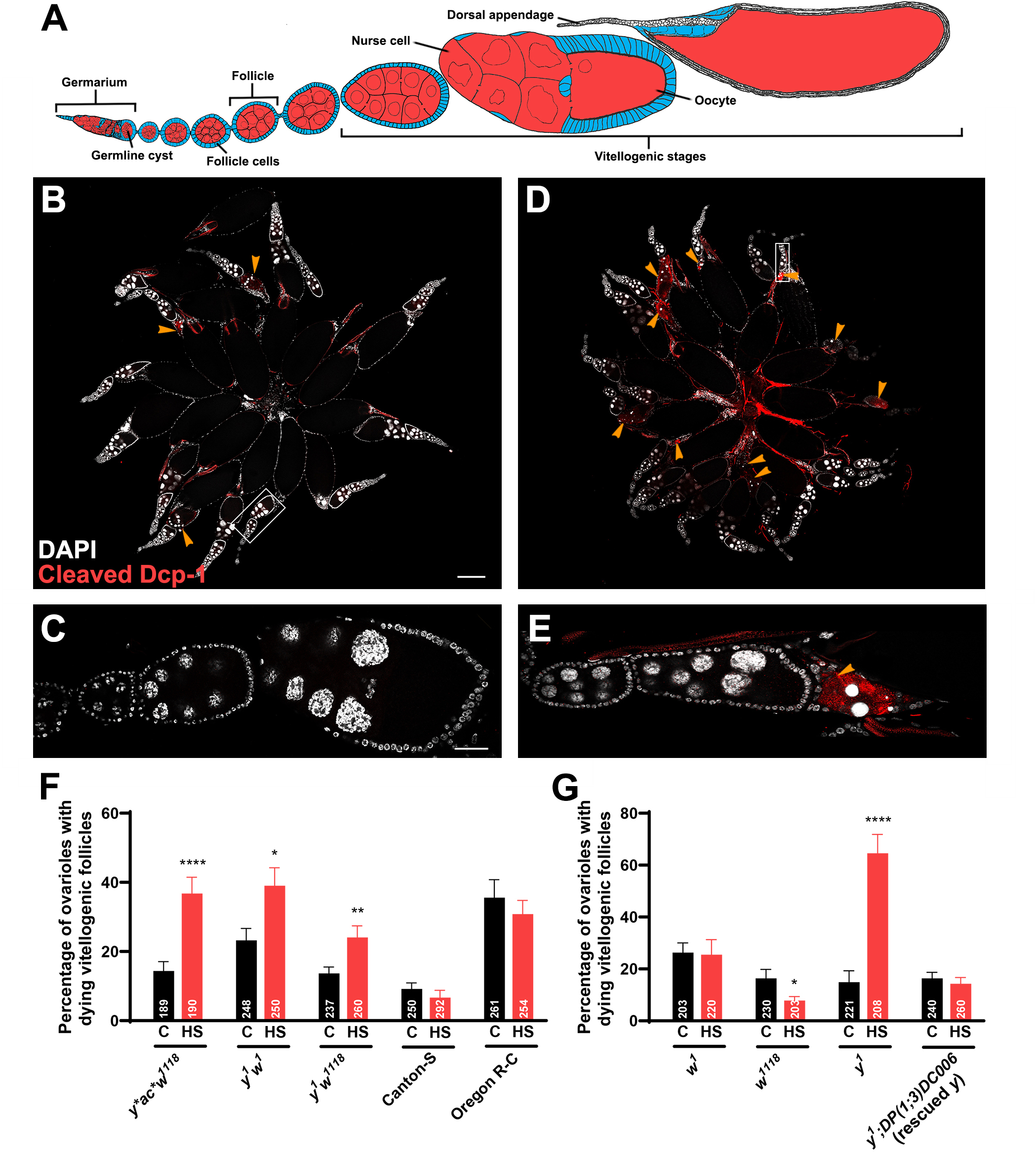
High sugar diet-induced vitellogenic follicle degeneration requires loss of *yellow* function. (A) *Drosophila* ovariole diagram showing the germarium followed by developing follicles (also known as egg chambers). Each follicle is composed of a 16-cell germline cyst (one oocyte and 15 supportive nurse cells) surrounded by follicle cells. Vitellogenic stages are characterized by the presence of yolk in the oocyte and mature oocytes have a fully formed dorsal appendage. (B–E) Single confocal sections of ovaries from *y* ac* w^1118^* females maintained on control (B,C) or high sugar (D,E) diets for 7 days. (C,E) Insets showing part of ovarioles from (B,D), respectively, with anterior to the left and posterior to the right. Cleaved Dcp-1 (red) labels apoptotic cells; DAPI (white) labels nuclei. (Anti-cleaved Dcp-1 antibodies also label dorsal appendages.) Dying vitellogenic follicles (indicated by arrowheads) are recognized by the presence of pyknotic nuclei and cleaved Dcp-1. Scale bars: 50 μM (B,D); 200 μM (D,E). (F,G) Frequencies of ovarioles containing dying vitellogenic follicles from females of the indicated genotypes maintained on control (black bars) or high sugar (red bars) diets. Numbers of ovarioles analyzed are shown inside bars. Data shown as mean±s.e.m. from two independent experiments. \**P*<0.05, \*\**P*<0.01, \*\*\*\**P*<0.0001; Chi-square test.

Genetic background, among other factors, influences diet-related phenotypes in organisms ranging from *Drosophila* to humans. For example, humans have high interpersonal variability in postprandial blood glucose levels (Zeevi et al. 2015); genetic predispositions influence the association between the consumption of sugar-sweetened beverages and adiposity (Haslam et al. 2018); 10-30% of obese individuals maintain healthy metabolic profiles (Schulze and Stefan 2024); and homozygosity of a specific variant of sucrase-isomaltase improves the metabolic health of adult cohorts in Greenland (Andersen et al. 2022). In mice, diet interacts with genetic background to affect weight gain and metabolic parameters (Reed et al. 2024), while in *Drosophila* genetic background affects the tolerance of developing larvae and pupae to high levels of dietary macronutrients (Havula et al. 2022).

*Drosophila* studies involving high sugar diets (Morris et al. 1999; Brookheart et al. 2017; Nunes and Drummond-Barbosa 2023) have routinely used flies carrying mutations in the *yellow* (*y*) and *white* (*w*) genes, which affect cuticle and eye pigmentation, respectively. *w* encodes an ABCG transporter, while *y* has unknown molecular function but is required for the production of melanin from dopamine during insect cuticle development (Mackenzie et al. 1999; Wittkopp et al. 2002; Drapeau 2003). *y* and *w* mutations are common in the *Drosophila* field because having them in the background facilitates the recognition of transgenes containing wildtype *y* and/or *w* “markers” (St Johnston 2013). It has remained unclear, however, whether the *y w* genetic background might affect how adult flies respond to high sugar diet, including their reduced female fertility phenotype.

In this study, we show that *y* mutation sensitizes females to a high sugar diet in a dopamine-dependent manner. Specifically, *y* mutant females have a higher frequency of vitellogenic follicle death on a high sugar diet, while wildtype Canton-S or Oregon-R-C females, or *y* mutants carrying a genomic rescue wildtype *y* transgene have no increase in follicle death in response to high sugar. Like *y* mutation, disruption of other genes in the melanin/sclerotin biosynthetic pathway (expected to alter dopamine levels) also leads to increased vitellogenic follicle death on a high sugar diet; conversely, alleviating dopamine imbalance in *y* mutant females prevents follicle death. Lack of dopamine production in the brain is sufficient to cause vitellogenic follicle death on high sugar, and this phenotype is partially reversed by dopamine dietary supplementation in adults. Finally, we show that more severe dopamine imbalance can cause follicle death regardless of diet or genetic background. These findings uncover a new role for *y* in sensitizing females to high sugar and an important connection between dopamine metabolism and oogenesis, providing a foundation for future research on the underlying cellular and molecular mechanisms. They also contribute more broadly to our understanding of how the effects of unhealthy diets differ depending on genetic factors.

## RESULTS

### The increase in vitellogenic follicle degeneration induced by a high sugar diet depends on genetic background

We previously showed that a high sugar diet, but not obesity, causes increased degeneration of vitellogenic follicles in *y* ac* w^1118^ Drosophila* females (Nunes and Drummond-Barbosa 2023). To determine if this high sugar effect depends on *y* and *w* mutations in the genetic background, we examined the ovarian response to a high sugar diet in *y^1^ w^1118^* and *y^1^ w^1^* females, in addition to two wildtype strains, Canton-S and Oregon-R-C (Öztürk-Çolak et al. 2024). Females raised on a standard control diet were maintained on either control or high sugar diets for 7 days and their ovaries analyzed using the cleaved Dcp1 apoptosis marker and pyknotic nuclei to identify dying follicles (Fig. 1B–E), as previously described (Armstrong and Drummond-Barbosa 2018; Nunes and Drummond-Barbosa 2023). As expected, a high sugar diet (compared to the control diet) led to increased death of vitellogenic follicles (Fig. 1F) in *y* ac* w^1118^* females. A similar response to high sugar was observed in *y^1^ w^1118^* and *y^1^ w^1^* females (Fig. 1F). By contrast, although wildtype Canton-S and Oregon-R-C females had different baseline levels of vitellogenic follicle death, a high sugar diet had no effect compared to a control diet in these females (Fig. 1F), showing that increased follicle degeneration in response to a high sugar diet depends on the genetic background.

### The increase in vitellogenic follicle degeneration induced by a high sugar diet requires loss of *yellow* function

We next tested *y* and *w* mutations individually for a role in sensitizing vitellogenic follicles to a high sugar diet. In homozygous null *w^1^* or strong hypomorphic *w^1118^* females, there was no increase in the percentage of ovarioles with dying vitellogenic follicles in response to a high sugar diet (Fig. 1G). By contrast, homozygous null *y^1^* females showed a significant increase in vitellogenic follicle death on a high sugar diet compared to a control diet, and this phenotype was rescued by a genomic rescue transgene carrying a wildtype copy of *y* (Fig. 1G). We conclude that mutation of *y* is specifically required for the increased death of vitellogenic follicles in response to a high sugar diet.

### Mutations in melanin/sclerotin pathway genes increase vitellogenic follicle degeneration on a high sugar diet

The *y* gene is required for the production of black melanin from dopamine, contributing to the pigmentation pattern of *Drosophila* adult cuticle and larval mouth parts (Nash and Yarkin 1974; Walter et al. 1996). Although the molecular function of Yellow remains largely unknown, the melanin biosynthesis pathway is well characterized in *Drosophila* (Massey and Wittkopp 2016; Sugumaran and Barek 2016) (Fig. 2A). Initially, tyrosine is converted to 3,4-dihydroxyphenylalanine (Dopa) by tyrosine hydroxylase (encoded by *pale* [*ple*]), and Dopa is decarboxylated into dopamine by Dopa decarboxylase (encoded by *Ddc*). Dopamine can then be converted into different types of pigments. For example, extracellular phenol oxidases (such as those encoded by *Prophenoloxidase 1* [*PPO1*], *PPO2*, and *PPO3*) generate black melanin (as part of a *yellow*-dependent pathway) and brown melanin. Dopamine can also be linked to β-alanine by the product of *ebony* (*e*) to form β-alanyl dopamine (NBAD), which is then polymerized into yellow-tan sclerotin by phenol oxidases; NBAD can also be converted back into dopamine by the hydrolase encoded by *tan* (*t*). Phenol oxidases also contribute to the production of the colorless polymer N-acetyl dopamine sclerotin from dopamine, and Ddc has been proposed to produce black melanin directly from Dopa (Massey and Wittkopp 2016; Sugumaran and Barek 2016).

**Fig. 2.**
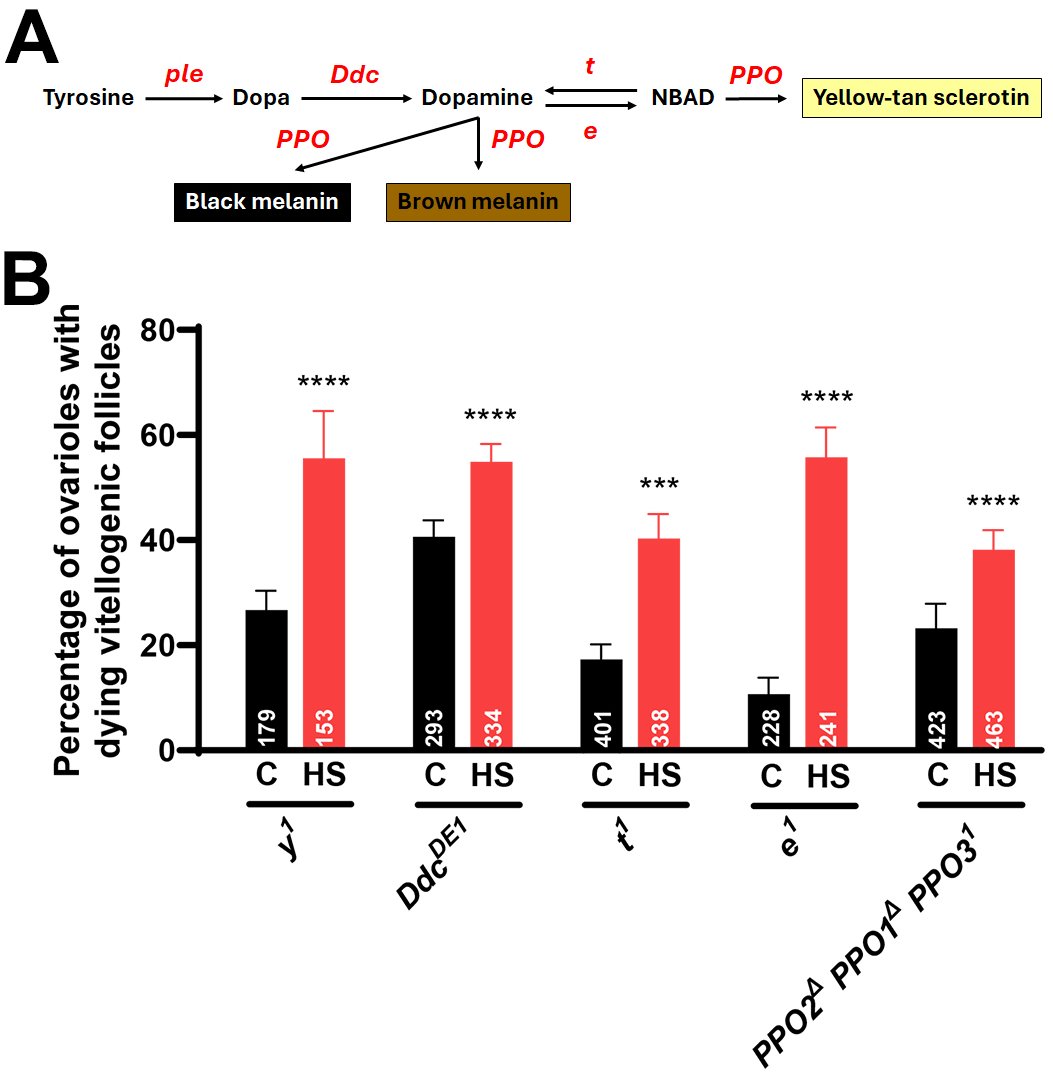
Mutations in melanin/sclerotin pathway genes increase vitellogenic follicle degeneration on a high sugar diet. (A) Simplified diagram of the melanin/sclerotin pathway (see text for details). (B) Frequencies of ovarioles containing dying vitellogenic follicles from females of the indicated genotypes maintained on control (black bars) or high sugar (red bars) diets. Numbers of ovarioles analyzed are shown inside bars. Data shown as mean±s.e.m. from two independent experiments. \*\*\**P*<0.001, \*\*\*\**P*<0.0001; Chi-square test.

To investigate if mutations in other components of the melanin/sclerotin biosynthesis pathway can lead to death of vitellogenic follicles on a high sugar diet, we tested females homozygous for hypomorphic *Ddc^DE1^*, *t^1^*, or *e^1^* alleles, as well as triple *PPO2^Δ^ PPO1^Δ^ PPO3^1^*homozygous females. As expected, *y^1^* mutant females showed increased death of vitellogenic follicles in response to a high sugar diet (Fig. 2B). Although the baseline frequency of ovarioles showing vitellogenic follicle death on the control diet differed across genotypes, in all cases there was a significant increase in vitellogenic follicle death on a high sugar diet (Fig. 2B). Considering the role of pigmentation genes in dopamine metabolism (Walter et al. 1996; Takahashi 2013), these results raised the possibility that changes in dopamine levels might contribute to the increased death of vitellogenic follicles on a high sugar diet in these mutant females.

### Decreased dopamine levels increase vitellogenic follicle death in *y^1^* females on a high sugar diet

In mammals, intermittent sugar intake causes brain dopamine release (Avena et al. 2008; Jacques et al. 2019); however, other studies have shown that long-term sugar consumption and obesity both result in low basal levels of dopamine in specific regions of the brain (Wang et al. 2001; Rada et al. 2005; Avena et al. 2008; Kravitz et al. 2016; Jacques et al. 2019; Wallace and Fordahl 2022). Consistent with these findings, male fruit flies fed a high sugar diet have decreased dopaminergic neuronal responses (May et al. 2020). We therefore hypothesized that maintaining females on a high sugar diet might lower their dopamine levels, leading to interactions with mutations in melanin/sclerotin pathway genes.

To test this hypothesis, we asked if dietary dopamine supplementation was sufficient to suppress the negative effects of a high sugar diet on the ovaries of *y^1^* females (Fig. 3). *y^1^* females on a high sugar diet supplemented with 5 mg/mL dopamine had a significantly lower frequency of ovarioles containing dying vitellogenic follicles (∼15%) than *y^1^* females on a high sugar diet alone (∼40%). Conversely, in *y^1^* females on a high sugar diet containing a tyrosine hydroxylase inhibitor, 3-Iodo-L-tyrosine (3IY; 1.25 mg/mL) to inhibit dopamine production (Cichewicz et al. 2017), the frequency of ovarioles with dying vitellogenic follicles remained elevated (Fig. 3). These results suggest that low dopamine levels and/or signaling underlie the effects of a high sugar diet in *y^1^* females. Considering the role of *y* in the synthesis of melanin from dopamine (Fig. 2A)—which might at first glance suggest that *y^1^* mutants would have higher levels of dopamine—our results also suggest a more complex role for *yellow* in dopamine metabolism and/or signaling.

**Fig. 3.**
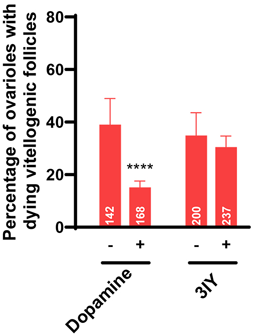
Dopamine supplementation suppresses vitellogenic follicle death in *y^1^* females on a high sugar diet. Frequencies of ovarioles containing dying vitellogenic follicles from *y^1^* females maintained on a high sugar diet with (+) or without (-) dopamine (5 mg/mL) or 3IY (1.25 mg/mL) supplementation. Numbers of ovarioles analyzed are shown inside bars. Data shown as mean±s.e.m. from two independent experiments. \*\*\*\**P*<0.0001; Chi-square test.

### Brain dopamine production is required to protect follicles from death on a high sugar diet

In addition to being produced in hypodermal cells during cuticle formation (Massey and Wittkopp 2016; Sugumaran and Barek 2016), dopamine is also produced in *Drosophila* brain neurons (Karam et al. 2020). We therefore asked whether brain dopamine production is important for the survival of vitellogenic follicles on a high sugar diet, taking advantage of an established brain dopamine-deficient *Drosophila* model (Cichewicz et al. 2017). In this model, homozygous *ple^2^*null females carry either of two genomic rescue transgenes. In control females, the genomic rescue transgene has a wildtype copy of *ple*, which restores expression of all tyrosine hydroxylase isoforms (Fig. 4A). Brain dopamine-deficient females have a modified genomic rescue transgene in which frameshift point mutations in *ple* specifically disrupt the translation of brain-specific isoforms of tyrosine hydroxylase while preserving the hypodermal isoform (Fig. 4A). While control females on a high sugar diet had low levels of dying vitellogenic follicles (Fig. 4B), brain dopamine-deficient females had a significant increase in the frequency of ovarioles with degenerating vitellogenic follicles (Fig. 4B). These results indicate that brain dopamine production is important for the survival of vitellogenic follicles. These results are consistent with the observation that *e^1^*mutants have increased vitellogenic follicle death on a high sugar diet (Fig. 2B), considering that loss of *e* function in glial cells has been shown to reduce dopamine production by brain neurons (Pantalia et al. 2023).

**Fig. 4.**
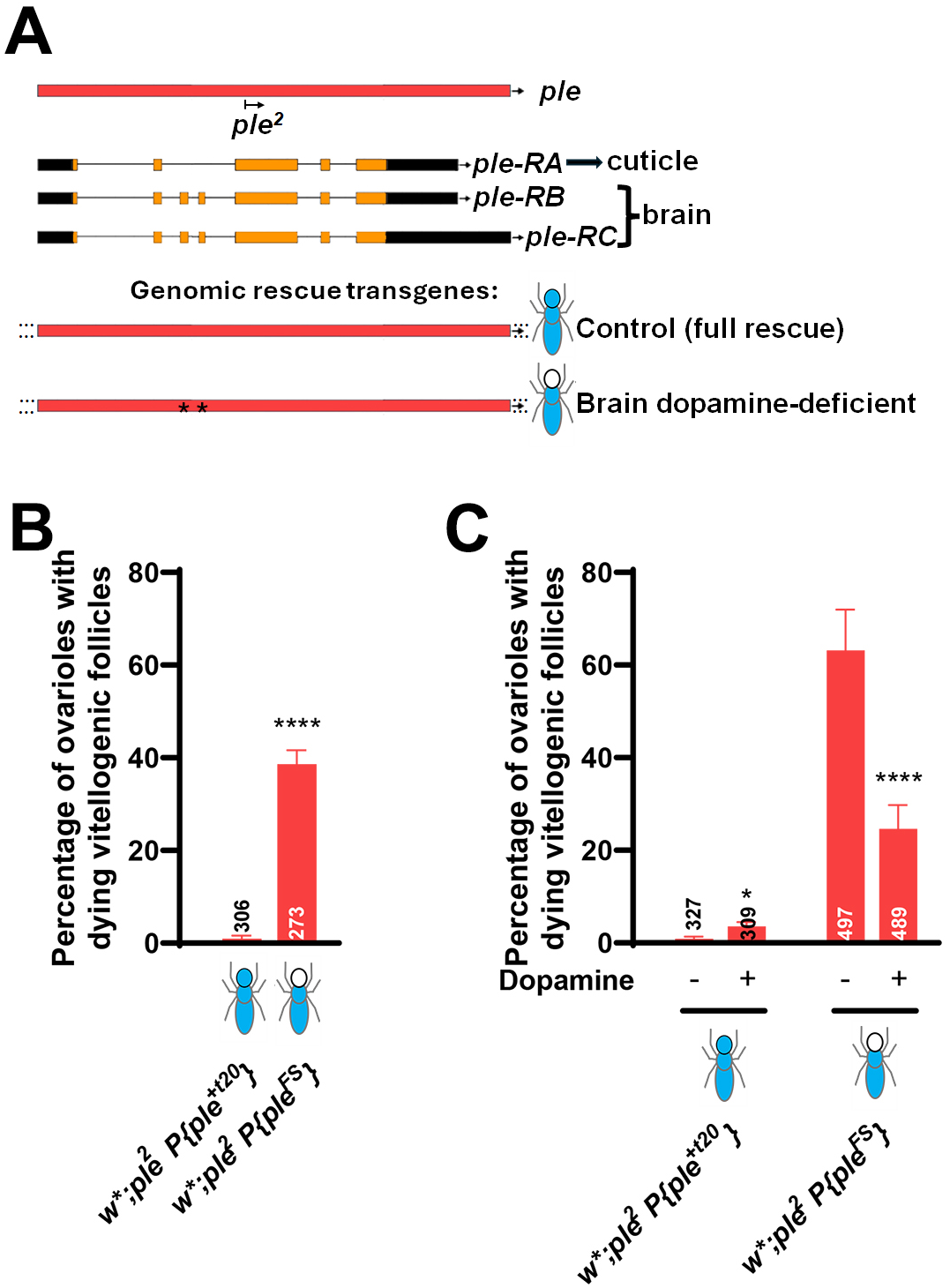
Brain dopamine protects vitellogenic follicles from death on a high sugar diet. (A) Diagram of brain dopamine-deficient flies and controls. The gene *ple* encodes three mRNA isoforms: *ple-RA* is expressed in hypodermal cells involved in cuticle formation, while *ple-RB* and *ple-RC* are expressed in the central nervous system. Females homozygous for the null *ple^2^* allele and carrying the wildtype genomic rescue transgene *P{ple^+t20^}attP2* are fully rescued (control), while *ple^2^* homozygous carrying a similar transgene with mutations specifically disrupting the brain isoforms, *P{ple^FS^}attP2*, only express the hypodermal isoform (brain-dopamine deficient) (Cichewicz et al. 2017). (B,C) Frequencies of ovarioles containing dying vitellogenic follicles from females of the indicated genotypes maintained on a high sugar diet for 7 days. In (C), females were maintained on a high sugar diet with (+) or without (-) dopamine (5 mg/mL) supplementation. Numbers of ovarioles analyzed are shown inside and/or above bars. Data shown as mean±s.e.m. from two independent experiments. \**P*<0.05, \*\*\*\**P*<0.0001; Chi-square test.

Interestingly, although dopamine is not thought to cross the blood–brain barrier in *Drosophila* (Cichewicz et al. 2017) or mammals (Calne et al. 1969; Cotzias et al. 1969), dopamine dietary supplementation partially suppressed the increased death of vitellogenic follicles in brain dopamine-deficient females (Fig. 4C). Based on these findings, we speculate that dopamine produced in the brain is released into the hemolymph as a neurohormone to modulate vitellogenic follicle survival. In support of this conjecture, previous studies have shown that neuronal-specific manipulations can drastically alter the levels of dopamine present in *Drosophila* hemolymph and mammalian blood (Goldstein and Holmes 2008; Zhao et al. 2010).

### Severe dopamine imbalance causes follicle death regardless of diet or genetic background

Our results indicated that dopamine balance is important for egg chamber survival on a high sugar diet (see Figs 3, 4), so we next investigated if this was also the case on a control diet and/or for wildtype females. We first tested if dopamine excess or deficit would increase follicle death in wildtype Canton-S females on a high sugar diet supplemented with dopamine (5 mg/mL) or 3IY (1.25 mg/mL). However, there were no significant effects at these concentrations (Fig. 5A). We therefore tested higher doses of dopamine (20 mg/mL) or 3IY (3 mg/mL). Canton-S females on a high sugar diet, where endogenous dopamine levels are presumably lower, were not sensitive to dietary dopamine but had elevated levels of vitellogenic follicle death upon inhibition of dopamine synthesis, indicating sensitivity to more severe decreases in dopamine levels (Fig. 5B). On a control diet, where endogenous dopamine levels are presumably higher than on high sugar, Canton-S females became sensitive to high dopamine supplementation and were also sensitive to high doses of 3IY (Fig. 5B). We also tested these pharmacological manipulations in wildtype Oregon-R-C females, which have higher basal levels of follicle death on both diets relative to Canton-S (Fig. 1F). In Oregon-R-C females, dopamine supplementation reduced vitellogenic follicle death on a high sugar diet, while inhibition of dopamine synthesis had no effect (Fig. 5C), suggesting that Oregon-R-C females might have lower levels of dopamine than Canton-S on a high sugar diet (Fig. 1F). On a control diet, Oregon-R-C females had increased levels of follicle death in response to either high dopamine supplementation or strong inhibition of dopamine synthesis (Fig. 5C), although it is unclear why the effects of 3IY were stronger on the control compared to high sugar diet. Our results suggest that dopamine levels and other interacting genes might differ among wildtype *Drosophila* strains, in agreement with previous behavioral studies (Colomb and Brembs 2014; Malacrida et al. 2022; Chaudhary et al. 2025). Nevertheless, based on our combined results, we conclude that, while *y* mutation sensitizes females to reduced dopamine levels caused by a high sugar diet (see Fig. 3), more severe dopamine imbalance— caused by brain dopamine deficiency (Fig. 4) or higher dopamine or 3IY dosage (Fig. 5)—is deleterious to vitellogenic follicles regardless of diet or genetic background. This apparent need for optimal levels of dopamine and/or dopamine signaling for oogenesis is consistent with other studies showing that fine tuning of dopamine levels is important in other contexts and organisms (Yamamoto and Seto 2014; Dobryakova et al. 2015; Zhou et al. 2023).

**Fig. 5.**
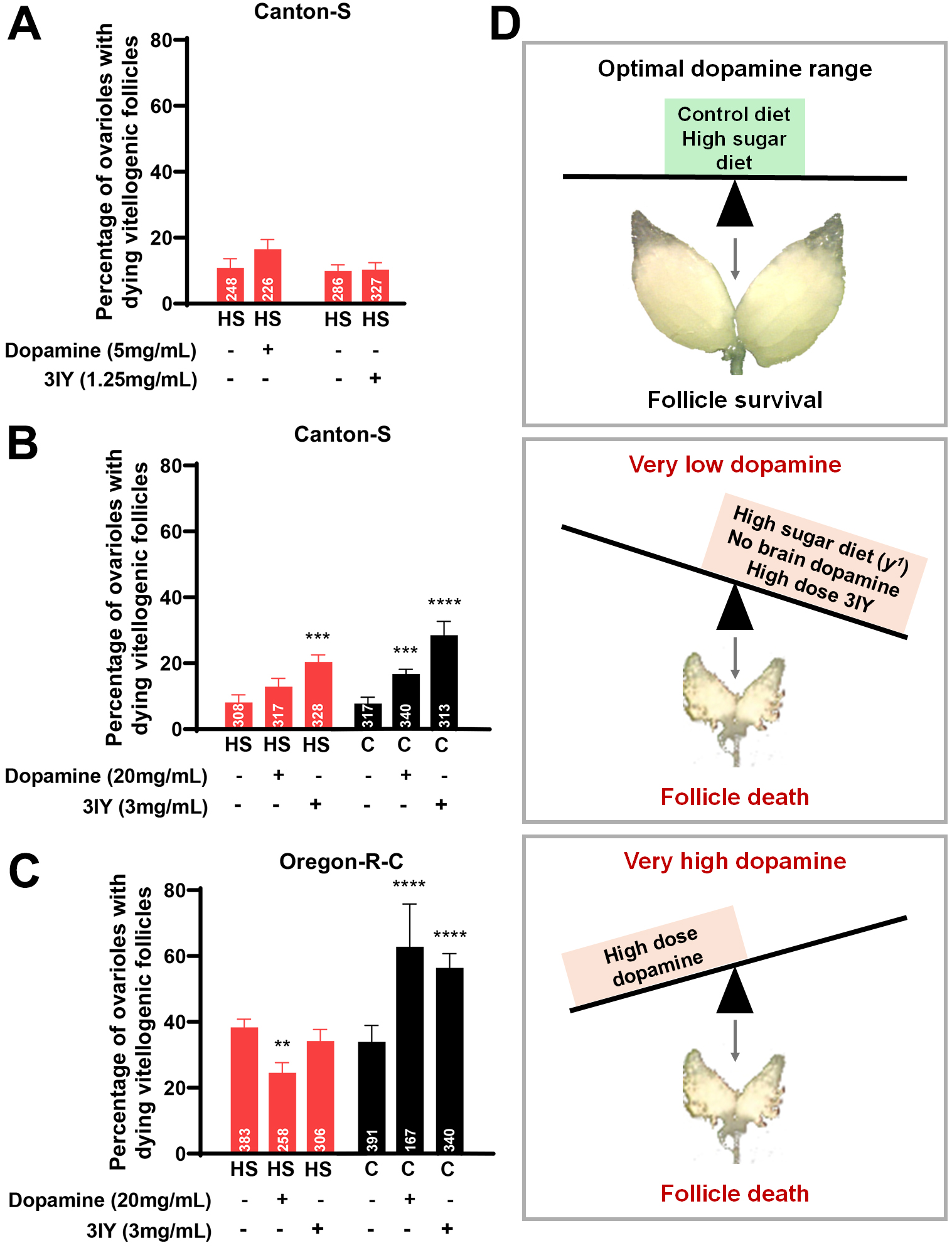
Dopamine imbalance causes follicle death regardless of diet or genetic background. (A–C) Frequencies of ovarioles containing dying vitellogenic follicles from Canton-S (A,B) or Oregon-R-C (C) females maintained on high sugar (red bars) or control (black bars) diets with or without dopamine [5 mg/mL in (A); 20 mg/mL in (B,C)] or 3IY [1.25 mg/mL] in (A); 3 mg/mL in (B,C)] supplementation for 7 days. Numbers of ovarioles analyzed are shown inside bars. Data are shown as mean±s.e.m. from two independent experiments. \*\**P*<0.01, \*\*\**P*<0.001, \*\*\*\**P*<0.0001; Chi-square test. **P<0.01. (D) Working model for dopamine imbalance in different contexts causing vitellogenic follicle death. Dopamine levels in the homeostatic range (*top*; e.g., in wildtype females regardless of diet; in *y^1^* females on the control diet) promote vitellogenic follicle survival. The frequency of vitellogenic follicle degeneration increases significantly when dopamine levels fall below (*middle*; e.g., in *y^1^* females on a high sugar diet, in brain dopamine-deficient females, or in wildtype females treated with a high dose of 3IY in specific diets) or above (*bottom*; e.g., wildtype females treated with a high dose of dopamine in specific diets).

## DISCUSSION

This study expands our understanding of how genetic background can influence responses to high sugar diets, focusing on the previously described ovarian response to a high sugar diet in *Drosophila melanogaster* (Nunes and Drummond-Barbosa 2023). Importantly, we uncover a role for brain-derived dopamine in modulating the survival of vitellogenic follicles in *Drosophila*. Based on our results, we propose the following working model (Fig. 5D): 1) An optimal range of dopamine levels contributes to the survival of vitellogenic follicles on a control diet. 2) Exposure of wildtype females to a high sugar diet leads to a relatively modest decrease in dopamine levels such that they remain within the optimal range; by contrast, *y* mutant females are more sensitive to a high sugar diet and dopamine therefore falls below the optimal range, causing increased follicle death. 3) In wildtype females on either a control or high sugar diet, increased follicle death requires more drastic alterations in dopamine levels induced either by genetic impairment of brain dopamine production or by high dose pharmacological manipulations that bring dopamine levels outside of the optimal range (i.e., either very high or very low). Our findings are consistent with observations that chronic consumption of high sugar is associated with decreased dopamine levels in mammals (Rada et al. 2005; Avena et al. 2008; Jacques et al. 2019). The vitellogenic follicle degeneration in *Drosophila* is also reminiscent of the increased follicle atresia observed in rats on high fat-high sugar diets (de Melo et al. 2021; Rejani et al. 2022; Demirel et al. 2024). Our study therefore raises the question of whether altered dopamine levels interact with diet and genetic background in other organisms, including humans, as a potential source of fertility issues or other chronic illnesses. This question is especially timely considering the prevalence of high sugar consumption in modern societies.

### Mechanisms underlying potentially conserved roles of dopamine in oogenesis?

Our findings align with previous research pointing to potential connections between dopamine and female reproduction in a wide range of organisms. For example, earlier studies showed that forcing either very low or very high levels of dopamine in newly eclosed Canton-S flies leads to reduced ovarian size (Neckameyer 1996), and cocaine, which interferes with dopamine signaling (Garris and Wightman 1995), increases vitellogenic follicle death (Sedore Willard et al. 2006). Inhibition of tyrosine hydroxylase in virgin fire ant *Solenopsis invicta* females lowers egg production (Boulay et al. 2001), while dietary dopamine supplementation was reported to increase ovary activation in queenless *Apis mellifera* workers (Dombroski et al. 2003). In the bumble bee *Bombus ignatis*, the brains of newly emerged queens have higher levels of dopamine compared to worker brains, likely driven by nutritional differences (Sasaki et al. 2021).

Interestingly, in *Drosophila sechellia*, which is a specialist fruit fly species adapted solely to the fruit of *Morinda citrifolia*, specific polymorphisms cause dopamine deficiency and oogenesis arrest unless these flies ingest L-DOPA present in *Morinda* (Lavista-Llanos et al. 2014). In the zebrafish *Danio rerio*, dopamine has an inhibitory role in female reproduction by antagonizing induction of pituitary follicle-stimulating hormone and luteinizing hormone by GnRH (Fontaine et al. 2013). Finally, in infertile women with hyperprolactinemia, long-term use of dopaminergic drugs can improve fertility (Crosignani 2012).

Other stresses and inputs besides high sugar also alter dopamine levels during different organismal responses. Dopamine neurons in the brain show strong responses to rewarding and aversive stimuli across organisms (Bromberg-Martin et al. 2010; Rosikon et al. 2023). Stresses including starvation, oxidative stress, and mechanical stress alter tyrosine hydroxylase activity levels in *Drosophila* and, conversely, reduction in tyrosine hydroxylase activity alters behavior and locomotive responses to stress (Neckameyer and Weinstein 2005). However, much remains unknown regarding the regulation and role of dopamine in the context of female fertility.

Our study opens new questions for future investigation regarding the role of dopamine in modulating *Drosophila* oogenesis. First, the *Drosophila* brain contains ∼50 morphologically distinct types of dopaminergic neurons organized in multiple clusters that innervate different regions of the brain and have different functions (Hartenstein et al. 2017; Siju et al. 2021; Marquis and Wilson 2022). What neurons serve as the key sources of dopamine in the control of oogenesis and how are they modulated by a high sugar diet? Second, *Drosophila* insulin-producing neurons have projections that extend outside of the brain to release insulin-like peptides directly into the hemolymph as neurohormones (Rulifson et al. 2002) that controls different processes during oogenesis (LaFever and Drummond-Barbosa 2005). Although dopaminergic neurons typically release dopamine as a neurotransmitter (for signaling to other neurons), there is evidence in flies and mammals suggesting that dopamine may also act as a neurohormone (Goldstein and Holmes 2008; Zhao et al. 2010). Are key dopaminergic neurons signaling to other neurons or releasing dopamine directly into the hemolymph to regulate follicle survival? Third, dopamine can act through different G protein-coupled receptors in two major classes (D1 and D2). *Drosophila* encodes four dopamine receptors including Dop1R1, Dop1R2, Dop2R, and the non-canonical receptor DopEcR (Siju et al. 2021). What dopamine receptors (and in which neurons/tissues) and downstream molecular, cellular, and physiological mechanisms are required for the effects of dopamine on oogenesis?

### Crosstalk between dopamine and serotonin during oogenesis?

In *Drosophila* and mammals, dopaminergic and serotonergic systems are known to interact with each other during development and in adults (Seo et al. 2008; Yarali et al. 2009; Neckameyer and Bhatt 2012; Niens et al. 2017; Kasture et al. 2018). For example, dopamine levels can affect serotonergic innervation in the brain and gut during *Drosophila* development (Neckameyer and Bhatt 2012; Niens et al. 2017). Dopamine and serotonin functionally interact in *Drosophila* and mammals to control different aspects of brain function, including long-term memory formation and behavior, and dopamine–serotonin imbalances are thought to contribute to depression, schizophrenia, and Parkinson’s disease (Seo et al. 2008). Serotonin has also been implicated in the control of oogenesis in the worm *Caenorhabditis elegans*, frogs, and mammals (Alhajeri et al. 2022; Aprison et al. 2022; Shmukler et al. 2022). The dopamine and serotonin biosynthetic pathways also have biochemical overlaps. Specifically, the DOPA decarboxylase encoded by *Ddc*, in addition to converting DOPA to dopamine (Fig. 2A), also decarboxylates 5-hydroxytryptophan to serotonin depending on neuronal type (Lundell and Hirsh 1994). Tetrahydrobiopterin is a cofactor for DOPA decarboxylase and for tryptophan phenylalanine hydroxylase (encoded by *henna*), which catalyzes the conversion of phenylalanine to tyrosine and of tryptophan to 5-hydroxytryptophan depending on substrate availability (Coleman and Neckameyer 2004). In our experiments, *Ddc^DE1^* hypomorphic mutants appeared to have elevated levels of vitellogenic follicle death (relative to *y^1^* and other pigmentation mutants; see Fig. 2) on a control diet. Although more severe alterations in dopamine levels in *Ddc^DE1^* females might explain this difference, we cannot exclude a possible contribution from altered serotonin levels. Future studies should more directly address whether dopamine functionally interacts with serotonin in the control of oogenesis.

### A connection between dietary water, high sugar, and dopamine?

We previously showed that dietary water supplementation can prevent the increased vitellogenic follicle death observed in *y* ac* w^1118^*females on a high sugar diet (Nunes and Drummond-Barbosa 2023). Interestingly, water can stimulate dopamine-producing neurons in both *Drosophila* and mammals (Lin et al. 2014; Jaksic et al. 2020; Mietlicki-Baase et al. 2021), and a subset of water-responsive neurons in *Drosophila* has also been shown to respond to sweet stimuli (May et al. 2020). It is tempting to speculate that water might reverse the effects of a high sugar diet on vitellogenic follicles by restoring dopamine homeostasis, although more research will be required to test this possibility.

### Yellow: the mystery remains

The *y* gene family is present across insect species (Ferguson et al. 2011; Gudelly et al. 2024). The *y* gene is best known for its requirement in the production of melanin from dopamine during cuticle development and there is intense research on its transcriptional and post-transcriptional regulation, especially in the context of the evolution of insect pigmentation (Wittkopp and Beldade 2009; Gudelly et al. 2024). The Yellow protein contains a Major Royal Jelly domain of unknown function and lacks enzymatic activity (Han et al. 2002; Claycomb et al. 2004; Drapeau et al. 2006; Ferguson et al. 2011). Although Yellow has been proposed to function as a structural scaffold (for pigment deposits or pigmentation enzymes) (Hinaux et al. 2018) or as a hormone (Drapeau 2003), its molecular function remains unknown. In this study we find a new connection between *y* function, sensitivity to dopamine imbalance in females on a high sugar diet, and follicle survival during *Drosophila* oogenesis, further highlighting the need to understand the molecular connection between Yellow and dopamine-related metabolism. Intriguingly, purified recombinant Yellow proteins from the sand fly *Lutzomya longipalpis* bind with relatively high affinity to dopamine and serotonin (Xu et al. 2011), adding a potential role in modulating the availability and/or stability of dopamine to the range of possibilities for how Yellow may act at the molecular level.

## MATERIALS AND METHODS

### *Drosophila* strains and culture conditions

*Drosophila* stocks were maintained at 22°C on standard medium composed of 5.8% v/v unsulfured cane molasses (Sweet Harvest Feeds), 4.64% w/v yellow cornmeal (Quaker), 1.74% w/v active dry yeast (Red Star), 0.93% w/v agar (Apex BioResearch Products), 1.05% Tegosept (Apex Chemicals) and 0.36% propionic acid (Apex Chemicals). (Note: The concentration of 5.8% molasses in our standard medium corresponds to ∼5% sugar, mostly sucrose.) The following strains are described in Flybase (Öztürk-Çolak et al. 2024): Canton-S (BDSC:64349), Oregon-R-C (BDSC:5), *y* ac* w^1118^* (Spradling 1986; Nunes and Drummond-Barbosa 2023), *y^1^ w^1^*(BDSC:1495), *y^1^ w^1118^*(BDSC:6598), *w^1^* (BDSC:145), *w^1118^* (BDSC:3605), *Ddc^DE1^* (BDSC:3168), *PPO2^Δ^ PPO1^Δ^ PPO3^1^*(BDSC:68387) *e^1^* (BDSC:1658), and *t^1^* (BDSC:130). *y^1^* (BDSC:169) is a null allele (Geyer et al. 1990), and the BAC transgene *DP(1;3)DC006* (BDSC:30217) (Venken et al. 2010) contains a wildtype copy of *y* (Öztürk-Çolak et al. 2024). *w*;ple^2^ P{ple^+t20^}attP2* and *w*;ple^2^ P{ple^FS^}attP2* (kind gifts from Jay Hirsch) have been previously described: *ple^2^* is a null allele; *P{ple^+t20^}attP2* is a 20 kb genomic rescue transgene encompassing *ple* that fully rescues *ple^2^*; and *P{ple^FS^}attP2* is a similar rescue transgene in which two compensating frameshift point mutations disrupt the translation of brain-specific *ple* isoforms but not of the hypodermal isoform for generation of brain dopamine-deficient flies (Cichewicz et al. 2017) (see Fig. 4A). For all experiments, newly eclosed flies raised on standard medium at 22°C were incubated at 25°C with 70% humidity and 12:12 light-dark cycle either on standard control medium (control diet) or standard medium with added sucrose (∼32% sugar content; high sugar diet) for 7 days.

### *yellow* rescue experiments

For *y* rescue experiments, *y^1^* homozygous mutants with or without the genomic rescue transgene were generated from the same cross. First, *w^1118^;;DP(1;3)DC006* males were crossed to *y^1^* virgin females to generate *y^1^;;DP(1;3)DC006/+* males. Then, *y^1^;;DP(1;3)DC006/+* males were crossed to *y^1^* virgin females to generate *y^1^*females (recognizable by their “yellow” cuticle) and sibling *y^1^;;DP(1;3)DC006/+* females (“rescued *y*”, which had wildtype cuticle pigmentation).

### Drug-feeding experiments

For drug feeding experiments, dopamine (Dopamine hydrochloride H8502, Sigma) and 3IY (3-Iodo-L-tyrosine I8250, Sigma) were directly mixed into melted medium at the indicated final concentrations and kept protected from light at 4°C for up to 1 week prior to use, as previously described (Cichewicz et al. 2017).

### Tissue staining, “flower mounting,” and microscopy

Ovaries were dissected in Grace’s Insect Medium (BioWhittaker) without teasing ovarioles apart. Samples were fixed for 15 minutes at room temperature in fixing solution [5.3% paraformaldehyde (Ted Pella) in Grace’s medium] and then rinsed and washed three times for 15 minutes each in PBST [0.1% Triton X-100 in PBS (10 mM NaH2PO4/NaHPO4, 175 mM NaCl, pH 7.4)]. Samples were blocked for 3 hours at room temperature in blocking solution [5% normal goat serum (MP Biochemicals) plus 5% bovine serum albumin (Sigma-Aldrich) in PBST] and then incubated overnight at 4°C in 1:100 rabbit anti-cleaved Dcp-1 (Cleaved *Drosophila* Dcp-1 (Asp215) 9578, Cell Signaling) in blocking solution. Ovaries were washed and incubated with 1:400 goat anti-rabbit conjugated to Alexa Fluor 568 (A-11011, Molecular Probes/Invitrogen) at room temperature and protected from light, as described (Armstrong and Drummond-Barbosa 2018). Samples were washed (protected from light) and stored in Vectashield containing DAPI (4′,6-diamidino-2-phenylindole, a fluorescent stain specific to DNA) (Vector Laboratories) overnight. Whole ovaries (“flower mounting”) were partially teased to keep all ovarioles attached to the main oviduct of each ovary and mounted on glass slides with coverslips. Data were collected using a Zeiss AxioImager-A2 fluorescence microscope or Zeiss LSM900 confocal microscope.

### Quantification of dying vitellogenic follicles

All degenerating vitellogenic follicles have pyknotic nuclei and cleaved Dcp-1 staining, which are not present in healthy vitellogenic follicles. All experiments were performed in two independent replicates with ovaries from multiple females (using only one ovary per female) per genotype and/or condition for each replicate, and statistical analysis was performed using Chi-square test analysis (Supplementary Data File S1).

## ACKNOWLEDMENTS

We thank the Developmental Studies Hybridoma Bank for antibodies, and Jay Hirsh (University of Virginia) and the Bloomington Stock Center (National Institutes of Health P400D018537) for *Drosophila* stocks. We thank Mallory Spencer, Emily Wessel, Ana Gandara, and Alicia Williams for careful reading of the manuscript and helpful editing suggestions. This work was supported by National Institutes of Health (NIH) grant R35 GM140857 (D.D.-B.).

